# Anti-citrullinated protein antibodies with diverse specificities ameliorate collagen antibody-induced arthritis in a time-dependent manner

**DOI:** 10.1101/2023.03.08.531560

**Authors:** Alejandro M. Gomez, R. Camille Brewer, Jae-Seung Moon, Suman Acharya, Sarah Kongpachith, Qian Wang, Shaghayegh Jahanbani, Heidi H. Wong, Tobias V. Lanz, Zelda Z. Love, Gundula Min-Oo, Anita Niedziela-Majka, William H. Robinson

## Abstract

Anti-citrullinated protein antibodies (ACPAs) are a hallmark of rheumatoid arthritis (RA) and have long been considered to contribute to pathogenesis. In this study, we sequenced the plasmablast antibody repertoires of RA patients and functionally characterized their encoded ACPAs. Recombinantly expressed monoclonal ACPAs bound citrullinated autoantigens, as well as autocitrullinated peptidylarginine deiminase 4 (PAD4). Using the collagen antibody induced arthritis (CAIA) mouse model, we demonstrated that the recombinant ACPAs significantly reduced paw thickness and arthritis severity as compared to isotype-matched control antibodies. Treatment with recombinant ACPAs also significantly reduced bone erosions, synovitis, and cartilage damage in histologic analysis of paws. This amelioration was observed for all the ACPAs tested and was independent of citrullinated antigen specificities. Furthermore, disease amelioration was more prominent when ACPAs were injected at earlier stages of CAIA than at later phases of the model, implying that ACPAs’ anti-inflammatory effects were more preventative than therapeutic. This study highlights a potential protective role for ACPAs in RA.

## INTRODUCTION

Rheumatoid arthritis (RA) is an autoimmune synovitis characterized by anti-citrullinated protein antibodies (ACPAs)^1^. Citrullinated antigens arise from the post-translational conversion of peptidyl arginine to peptidyl citrulline by peptidyl arginine deiminase (PAD) enzymes^2^. ACPAs are a hallmark of RA, yet there is significant debate regarding the role of ACPAs in the pathogenesis of RA^3^.

Sequencing of RA patients’ antibody repertoires and expression of the encoded ACPAs revealed that some ACPAs exhibit binding to several citrullinated epitopes, while other ACPAs recognize specific citrullinated antigens^4, 5^. ACPAs are approximately 65% sensitive and 97% specific for the diagnosis of RA^6^, but are also found in small subsets of other autoimmune diseases like Sjogren’s^7^ and anti-neutrophil cytoplasmic antibody (ANCA)-associated vasculitis^8^, and in a small fraction of healthy individuals^9^. ACPAs were first described in 1964 and have since been both directly and indirectly implicated in the pathophysiology of RA and joint inflammation^10^. In RA, ACPAs are associated with a higher risk for bone erosion^11^ and extra-articular complications^12^. In addition, high ACPA titers and epitope spreading of the ACPA response have been associated with RA development in at-risk individuals^13, 14^, and with higher disease activity in established RA^15^.

The direct pathogenic role of ACPAs in joint inflammation, however, has been primarily investigated in animal models, with mixed results. In one line of studies, injection of pooled human ACPAs into mice induced pain-like behavior^16^, and induced osteopenia and increased osteoclastogenesis^17^. In another line of studies, murine ACPAs generated in response to citrullinated histone H2B did not induce arthritis in isolation but exacerbated inflammation in mice with low-level collagen-induced arthritis (CIA)^18^. Likewise, injection of an anti-citrullinated fibrinogen ACPA into animals did not induce joint inflammation, unless it was delivered in combination with sub-optimal doses of anti-collagen type II antibodies^19^. Altogether, these findings support a pathogenic role for ACPAs in arthritis. In contrast, it was recently reported that recombinant ACPAs ameliorated disease symptoms in collagen-antibody induced arthritis (CAIA)^20, 21^, and in CIA and various models of neutrophil extracellular trap-(NET) mediated diseases^22^, suggesting that ACPAs can be protective in arthritis.

In this study, we functionally characterized clonal ACPAs from RA. We showed that patient-derived ACPAs, as compared to isotype-matched control antibodies, reduced the severity of CAIA, regardless of their citrullinated antigen specificities. Interestingly, this amelioration was prominent at earlier stages of the model and was not observed at later stages of CAIA.

## MATERIALS AND METHODS

### RA patient’s antibody repertoire sequencing and generation of recombinant antibodies

PBMCs from rheumatoid arthritis (RA) patients were isolated and plasmablasts were sorted by flow cytometry, and subjected to paired heavy and light chain antibody repertoire sequencing at the single cell level^5^. Select plasmablast clonal family antibodies were gene synthesized and recombinantly expressed in a human IgG1 backbone, as previously described^5, 23^. Antibodies derived from healthy individuals targeting influenza^24^ and from neonatal lupus-associated congenital heart block subjects were selected as in-house produced negative controls. Commercially available human isotype control IgG1 (cat. 403502; BioLegend Inc.) was also used as a negative control. Recombinant antibodies for animal experiments were cloned in a mouse IgG2a backbone after codon optimizing the corresponding V(D)J regions using proprietary mHC and mLC immunoglobulin (Ig) plasmids (LakePharma/Curia Global Inc. and Gilead Sciences). Ultra-LEAF™ Purified Mouse IgG2a, κ was used as isotype control (cat. 400282; BioLegend). Immunoglobulin concentration was estimated by NanoDrop (ThermoFisher) or human IgG ELISAs (Bethyl Laboratories).

### Planar antigen arrays and ELISAs

Planar antigen arrays were printed, probed, and scanned as previously described^25, 26^. The arrays contained approximately 320 antigens per slide, primarily 19-mer peptides and some full proteins of RA-associated autoantigens. They included 51 citrullinated autoantigens and their native counterparts, plus additional native autoantigens and non-RA relevant proteins, to assess both the polyspecificity and the polyreactivity of tested antibodies^27^. We considered an antibody to be polyspecific when it bound more than 10 RA-autoantigens with mean fluorescent units (MFIs) higher than the mean MFI of the array and polyreactive when it bound to non-RA relevant proteins. Antibodies were probed at 50 μg/mL on array slides with quadruplicate print-spots/antigen. Slides were incubated for 1.5h at 4C, washed, and incubated with a 1:2,500 dilution of Cy3-conjugated goat anti-human IgG/IgM secondary antibody (Jackson ImmunoResearch), followed by washing and scanning of fluorescence intensities. Raw MFIs were background corrected by subtracting the mean+3xSD of all negative control antibodies (4/assay) for a given antigen to the corresponding MFIs for that antigen. Negative values were converted into 1 MFI. ELISAs were conducted with recombinant full proteins coated overnight in Nunc MaxiSorp 384 plates (ThermoFisher) (2µg/ml, 50µl/well) in bicarbonate/carbonate buffer (pH 9.5). Plates were blocked with 2% BSA in PBS for 1h at room temperature (RT). Antibodies were probed at 1µg/ml and incubated for 1h at RT. Antibody binding was detected with HRP-conjugated goat anti-human IgG Fc antibody (Bethyl Laboratories). Plates were washed five times with PBS-Tween between steps. After adding the developing reagent (Super AquaBlue Substrate, ThermoFisher), plates were monitored at OD405 with a plate reader (BioTek). The recording was terminated when positive controls reached ∼2.5. ODs were blank-corrected and the fold-over isotype control were calculated for comparisons between antigens.

PAD4 autoantibody ELISA Kits (cat. 500930; Cayman Chemical) were run according to manufacturer recommendations. Antibodies were tested at 1-10 μg/ml, in duplicate. Anti-PAD4 U/mL were inferred from the kit’s standard curve and converted into mg/mL of anti-PAD4 Ig with the conversion factor 1U = 1ng anti-PAD4 Ig. The cut-off value for positivity was defined as the mean + 3xSD of isotype control replicates.

### Bio-layer interferometry

The binding affinity of recombinant antibodies to antigens was estimated by bio-layer interferometry (Octet^®^ QK, ForteBio/Sartorius). Antibodies were loaded at 10-20 nM onto anti-human IgG Fc capture or anti-Mouse IgG Fc capture biosensors. Antigen targets were probed in serial dilutions ranging from 5000 to 15.6 nM in 1X kinetic buffer (ForteBio/Sartorius). Equilibrium dissociation constants (*K*_D_’s) were calculated from K_on_ and K_off_ kinetic parameters. The reported *K*_D_’s were computed from globally fitted binding curves (Langmuir 1:1 interaction model; between 2-8 curves/analyte), with the Octet^®^ BLI Analysis software (ForteBio/Sartorius). Measurements from blank wells and isotype control sensors were subtracted from values. Only curves that had at least 0.2 nm of blank-corrected response and >0.9 R^2^ in their local fit were included in the global fit calculations.

### Animal experiments

All animal studies were conducted following standard procedures approved by Stanford’s Administrative Panel on Laboratory Animal Care (APLAC; Protocol #9942). Mice were purchased from The Jackson Laboratory and housed in pathogen-free conditions (between 3-5 animals/cage), with ad libitum access to water and chow. Male C57BL/6 (#000664) and DBA/1J (#000670) mice between 7-8 weeks were used for experimentation.

The collagen antibody induced arthritis (CAIA) was induced by retro-orbital injection of a cocktail of 5 monoclonal antibodies (mAbs) against mouse collagen type II (anti-CII cocktail, Chondrex Inc), followed by intraperitoneal injection of lipopolysaccharide (LPS) (50 μg/mouse) three days later. For CAIA amelioration experiments, the optimal dose of anti-CII cocktail (5 mg/mouse for C57BL/6 and 1.5 mg/mouse for DBA/1J) was used to ensure maximal joint inflammation and incidence of arthritis. For evaluating if patient-derived mAbs could enhance joint inflammation, a sub-optimal form of CAIA was developed. The dose of anti-CII cocktail for inducing sub-optimal CAIA was optimized in-house and was defined as 2.5 mg/mouse for C57BL/6 and 1 mg/mouse for DBA/1J, as detailed in Suppl. Fig1.

Recombinant antibodies were injected retro-orbitally at different time-points, depending on the CAIA experiment (see Results section). When combined into treatment groups of 3 ACPAs, 0.5 mg/mAb/animal were mixed for a total of 1.5 mg mAb/animal/injection. Groups treated with a single mAb received 1.5 mg mAb/animal/injection. Recombinant antibodies were delivered in 20 mM Histidine, 9% Sucrose, 0.05% Polysorbate 80, pH 5.8 buffer, verified for low endotoxin content (<1 EU/mg). No-antibody CAIA controls were injected with buffer alone and wild-type (WT) naïve controls were left untreated. Mice were weighed and clinically scored every 2-3 days. Joint inflammation was scored by blinded investigators following recommendations from Chondrex Inc.: each limb was graded on a scale from 0-4, for a maximum of 16 per animal. In addition, paw thickness was measured with a flat-anvil thickness gauge (Mitutoyo 7300S) on days in which animals were clinically scored. Accumulated change in paw thickness was calculated as a summation of all changes (as a percentage of the previous time-point measurement) following the first antibody injection. Animals were sacrificed between 13-18 days post-CAIA induction by CO2 inhalation followed by cervical dislocation. Terminal blood was collected by retro-orbital bleeding for heparinized plasma separation. All limbs were harvested and either fixed in formalin for histology or flash-frozen for downstream analysis. Mice with severe arthritis (score of 3+ on all 4 paws) that experienced >20% weight loss, or became pre-morbid, were sacrificed before the experimental endpoint. Mice that failed to manifest any arthritis by day 7 post-induction, and did not receive any antibody treatment until then, were excluded. Total number of mice used in animal experiments: 153; number of mice excluded from analysis: 9.

### Histology

Formalin-fixed paws were embedded in paraffin, sectioned onto glass slides (2 sagittal-sections/slide), and stained with hematoxylin and eosin (Premier Laboratory). Slides were imaged in bright field with a 2× objective lens in a digital microscope (BZ-X700; Keyence) and scored for synovial inflammation, cartilage damage, and bone erosion by a blinded investigator (QW), adapting published guidelines^28^.

### Statistics and Graphics

GraphPad Prism 9 for macOS (Version 9.4.1; GraphPad Software) was used for generating graphs and for statistical analysis. Normal distribution was confirmed by normality tests for datasets that were analyzed by unpaired ANOVAs. Non-gaussian datasets were analyzed with unpaired Kruskal-Wallis tests. For animal studies, a mixed-effects model (if the dataset had missing values) or repeated-measures 2-way ANOVA (no missing values) were used for statistical analysis of variables quantified at multiple time-points. Turkey’s (parametric) and Dunn’s (non-parametric) tests were used for multiple comparisons. 2-sided *p* values: ns: p > 0.05; *: *p* ≤ 0.05; **: *p* ≤ 0.01; ***: *p* ≤ 0.001.

## RESULTS

### Clonal expansions of ACPAs encode antibodies polyspecific for citrullinated autoantigens, including PAD enzymes

We previously sequenced the plasmablast antibody repertoire of patients with RA^5^. In the present study, we expanded the antigen profiling and functional characterization of rheumatoid arthritis (RA) plasmablast-encoded anti-citrullinated protein antibodies (ACPAs) (Fig. 1A). We recombinantly expressed a set of 20 monoclonal antibodies (mAbs) which included all members of the three main ACPA clonal families (CF) of an RA patient, plus one singleton ACPA, and two non-ACPAs as control mAbs.

**Figure 1.**
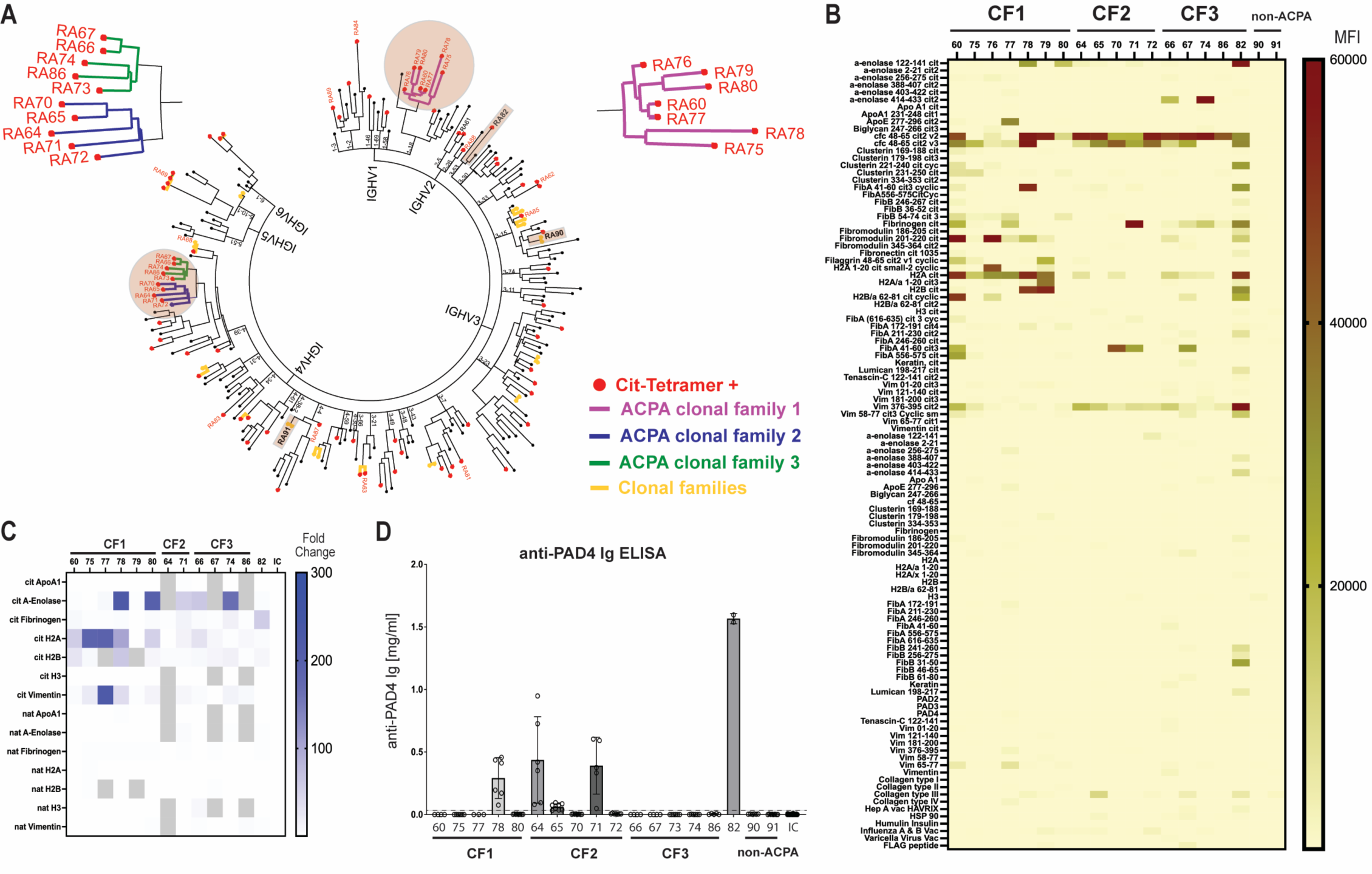
RA patient-derived, clonally expanded ACPAs target multiple citrullinated autoantigens, including citrullinated PAD4. **A.** Phylogenetic tree of the plasmablast immunoglobulin genes from a CCP+ RA patient with multiple ACPA clonal expansions. The most prominent ACPA clonal families are enlarged for better visualization. Red dots at branch tips indicate antibody clones that bound to citrullinated tetramers during FACS sorting. Antibodies belonging to a clonal family are denoted by highlighted, colored branches. Non-clonal ACPAs and non-ACPA controls that were expressed for downstream analysis are identified by shaded rectangles **B.** Binding of recombinant antibodies to RA-associated human autoantigens printed on planar arrays, in both their native and citrullinated forms. Non-RA relevant proteins (insulin, viral proteins, flag peptide) were included to assess polyreactivity. Antibodies from all ACPA clonal families were tested, plus one singleton and two non-ACPA controls. Raw mean-fluorescence intensities (MFIs) were background corrected by subtracting the mean+3xSD MFIs of 4 negative control antibodies for the corresponding antigen. The background-corrected MFIs of 4 replicate spots per antigen are shown. Only citrullinated antigens, their corresponding native counterparts, and non-RA relevant proteins are shown in the heatmap **C.** ACPAs reactivity to full protein, RA-associated human autoantigens on enzyme-linked immunosorbent assays (ELISAs). The heatmap summarizes ACPAs’ reactivity to recombinant citrullinated proteins and their native counterparts. Recombinant antibodies were tested at 1 μg/mL, in triplicate. Values displayed are folds over isotype control antibody signal (OD405) and they are the average of two independent experiments. Grey-shaded squares: not tested **D.** Targeting of PAD4 by patient-derived ACPAs. Recombinant antibodies were assayed on anti-PAD4 autoantibody ELISA kits (Cayman Chemical). Antibodies were tested at 1-10 μg/ml, in duplicate. Values shown are the calculated mg/mL of standardized anti-PAD4 antibody, presented as the mean +/− SD of at least two independent experiments. The dashed line represents the positivity cut-off threshold, defined as the mean+3xSD of isotype control replicates. *CF = clonal family; cf = citrullinated filaggrin; cit = citrullinated; cyc = cyclic; FibA/B = fibrinogen α/ β ; H2A/B, H3 = histone 2A/B, 3; Hep = hepatitis; HSP = heat shock protein; IC: isotype control; IGHV = immunoglobulin heavy-chain variable gene; nat = native; SD = standard deviation; Vac = vaccine; Vim = vimentin*

First, we characterized the autoantigen specificities of this set of recombinant mAbs by planar antigen arrays. The arrays contained 320 19-mer peptides and full proteins encompassing many autoantigens associated to RA, in both their native and citrullinated (cit) forms, as well as non-RA relevant antigens (insulin, viral proteins, flag peptide). The main citrullinated peptides targeted by recombinant ACPAs belonged to citVimentin, citAlpha-Enolase, citFibromodulin, citFibrinogen A, citHistones (H2A/H2B), and citClusterin (Fig. 1B). Very little to no reactivity was observed for the native peptide counterparts. Most ACPAs bound strongly to cyclic citrullinated fibrin peptides (cfc), the main component of early-generation commercial CCP assays, corroborating the ACPA classification of these mAbs. Certain ACPAs (e.g. RA60, RA78, RA82), detected multiple citrullinated peptides, from various proteins, suggesting polyspecificity for citrullinated antigens. ACPAs did not bind to non-RA relevant antigens, suggesting that they were not polyreactive (i.e. binding to multiple, unrelated antigens). Non-ACPA control mAbs did not bind any antigen on these arrays.

We then validated the main ACPA targets by ELISAs, using both native and citrullinated full-protein versions of the most reactive antigens from planar arrays. Representative mAbs from all ACPA clonal families bound preferentially to citrullinated proteins and not to their native counterparts (Fig. 1C) (mean fold change for binding to cit-over native antigens: 27.3). ACPAs’ reactivity on ELISAs corroborated most planar array targets, except for citVimentin and citFibrinogen, which were less reactive on ELISAs. Polyspecificity for citrullinated proteins was again observed in some ACPAs (RA77, RA78, RA80). In addition, we screened all recombinant mAbs for binding to PAD4 by ELISAs. This enzyme was of particular interest because of its association with citrullination and neutrophil extracellular traps (NETs)^29^. Previous reports suggested a pathogenic role for a subgroup of cross-reactive anti-PAD4/3 antibodies in RA^30^. Commercial ELISAs identified several ACPAs with reactivity to PAD4, especially mAbs from CF2 and polyspecific ACPAs (Fig. 1D). To assess cross-reactivity to PAD3 and binding to citrullinated forms of PAD enzymes, ACPAs were tested on in-house ELISAs coated with either native or auto-citrullinated forms of PAD4 or PAD3. As shown in Suppl. Fig2, ACPAs’ reactivity to PAD4 replicated the results from commercial ELISAs, with strong binding to the auto-citrullinated form of the enzyme. Several PAD4+ ACPAs also bound to PAD3 on ELISAs, though their preference for auto-citrullinated PAD3 was less pronounced than for citPAD4 (5.7-fold less, on average).

Further, we evaluated the binding affinity of recombinant ACPAs to several of their ELISA-corroborated antigens, both native and citrullinated, by biolayer interferometry (BLI) (Table 1). ACPAs bound citrullinated antigens with equilibrium dissociation constants (*K*_D_’s) ranging from 0.1 – 451 nM. *K*_D_’s were calculated for all ACPAs that bound PAD4 on ELISAs, confirming their high affinity for the enzyme. The majority of PAD4+ ACPAs bound only to autocitrullinated PAD4 on BLI, excepting most mAbs from CF2, which bound also to native PAD4, though with lower affinity (350-950 fold less). Binding to PAD3 was confirmed by BLI only for RA64 and RA65, which bound to the native enzyme alone. Most PAD+ ACPAs were restricted to CF2, along with some polyspecific ACPAs that targeted PAD4. Furthermore, we generated ancestral members of CF2 by reverting the affinity-matured antibodies back to their germline sequence with IgTree^31^. Several of these CF2 parental antibodies interacted with both native and citrullinated PAD4 on BLI (Suppl. Fig3). Their affinity for autocitrullinated PAD4 was stronger and more prevalent than for the native enzyme. In fact, most ancestral members of CF2, including the germline sequence, bound strongly to citPAD4, suggesting that this antigen was a main driver for affinity maturation within this ACPA clonal family. Interestingly, ACPAs in CF2 also bound PAD4 substrates (histones H3 and H4) with high affinity (*K*_D_’s in the nM range). They targeted both native and citrullinated histones, with a preference for citH4 and, unexpectedly, native H3. A similar pattern was observed in the other clonal families. High affinity binding to citFibrinogen was also validated for several ACPAs (*K*_D_’s between 0.1-6.6 nM), across all clonal families. No binding was observed for native fibrinogen, and no interaction with PAD2, vimentin, or alpha enolase was confirmed by BLI.

**Table 1.** Binding affinities of ACPAs to native and citrullinated RA-associated autoantigens. Equilibrium dissociation constants (K_D_’s) were determined by bio-layer interferometry (BLI) for select antibody/antigen pairs. K_D_’s are reported in nM and were calculated by globally fitting response curves obtained for at least 2 serially diluted antigen concentrations per analyzed ACPA. Measurements from blank wells and isotype control sensors were subtracted from values. Only curves that had at least 0.2 nm of blank-corrected BLI response and >0.9 R^2^ in their local fit were included in the global fit calculations. Zeros denote no binding and blanks indicate not tested. CF = clonal family; cit = citrullinated; mAb = monoclonal antibody; nat = native

| CF | mAb | PAD4 | | PAD2 | | PAD3 | | Histone H3 | | Histone H4 | | Fibrinogen | | Vimentin | | α-Enolase | |
| --- | --- | --- | --- | --- | --- | --- | --- | --- | --- | --- | --- | --- | --- | --- | --- | --- | --- |
|  |  | cit | nat | cit | nat | cit | nat | cit | nat | cit | nat | cit | nat | cit | nat | cit | nat |
| 1 | RA60 |  | 0 |  |  |  |  |  |  |  |  | 0 | 0 |  |  |  |  |
| 1 | RA75 | 0 | 0 |  |  | 0 |  | 0 | 0 |  |  | 0.8 | 0 | 0 | 0 | 0 | 0 |
| 1 | RA76 |  |  | 0 | 0 |  | 0 |  |  |  |  |  |  |  |  |  |  |
| 1 | RA77 |  |  |  |  |  |  |  |  |  |  |  |  |  |  |  |  |
| 1 | RA78 | 2 | 0 |  |  | 0 | 0 | 115 | 112 | 2.5 | 28 | 6.6 | 0 | 0 | 0 | 0 | 0 |
| 1 | RA79 |  |  |  |  |  |  |  |  |  |  |  |  |  |  |  |  |
| 1 | RA80 | 328 | 0 | 0 | 0 |  | 0 | 0 | 0 | 2.1 | 16 | 0 | 0 | 0 | 0 | 0 | 0 |
| 2 | RA66 | 0 | 0 | 0 | 0 |  |  | 21 |  | 0.6 | 16 | 0 | 0 | 0 | 0 | 0 | 0 |
| 2 | RA67 |  | 0 | 0 | 0 |  |  |  |  |  |  | 0 | 0 |  |  |  |  |
| 2 | RA73 | 0 | 0 |  |  |  | 0 | 359 | 1.7 |  |  | 0 | 0 |  |  |  |  |
| 2 | RA74 | 0 | 0 |  |  |  |  | 73 | 643 | 3.2 | 23 | 2.1 | 0 | 0 | 0 | 0 | 0 |
| 2 | RA86 | 0 | 0 | 0 |  |  | 0 |  |  |  |  |  |  | 0 | 0 |  |  |
| 3 | RA64 | 0.2 | 132 | 0 | 0 | 0 | 33 | 342 | 48 | 1.4 | 14 | 0 | 0 | 0 | 0 | 0 | 0 |
| 3 | RA65 | 1 | 996 |  |  | 0 | 20 | 451 | 34 | 2.5 | 25 | 0.15 | 0 | 0 | 0 | 0 | 0 |
| 3 | RA70 | 0 | 0 |  |  | 0 | 0 | 442 | 27 |  |  | 0 | 0 |  |  |  |  |
| 3 | RA71 | 1.4 | 499 | 0 |  | 0 | 0 | 0 | 463 | 0.3 | 26 | 0.1 | 0 | 0 | 0 | 0 | 0 |
| 3 | RA72 | 0.7 | 0 |  |  | 0 | 0 | 0 | 21 |  |  |  | 0 |  |  |  |  |
| - | RA82 | 0.4 | 0 |  |  | 0 | 0 | 0 | 0 | 8.2 | 0 | 1.2 | 0 | 0 | 0 | 0 | 0 |

Thus, we observed citrullinated-antigen reactivity patterns within clonal families, particularly against citPAD4. Binding to citPAD4 was quite prominent within CF2, all members bound to citPAD4 on ELISAs and *K*_D_’s were calculated for most of them. CF3, on the other hand, did not bind citPAD4 at all. citPAD4, however, was also targeted by the more polyspecific members of CF1 (Fig. 1, Table 1). Most clonal ACPAs also bound, to some extent, to native forms of certain RA-associated antigens, particularly to histones and PAD4. This was observed both on ELISAs and on BLI. Binding to the citrullinated form of these antigens, however, was stronger than for the native form (mean fold change cit/nat: 231). One exception was RA82, a non-clonal ACPA, for which only binding to the citrullinated form of the antigens was confirmed by BLI (Table 1).

In summary, ACPA clonal expansions bound preferentially to citrullinated autoantigens commonly associated to RA but were frequently polyspecific in their citrullinated antigen recognition. Targeting of citPAD4 and histone H3 was more restricted to CF2, while histones H3-H4 and fibrinogen were the main citrullinated targets in CF1 and CF3. Polyspecificity was more evident in ELISAs and protein arrays, where a wider selection of citrullinated autoantigens was detected by clonal ACPAs.

### Recombinant ACPAs with diverse citrullinated-antigen specificities ameliorate CAIA

Recombinant ACPAs were further tested for their functional effects in a collagen-antibody-induced arthritis (CAIA) model in C57BL/6 mice. ACPAs were subdivided into treatment groups based on their clonality and specificities, which included: citPAD4+/CF2 ACPAs (RA64, RA65, RA71), PAD4-/CF1-3 ACPAs (RA66, RA74, RA75), and polyspecific ACPAs (RA78, RA80, RA82). For animal experiments, human ACPAs were expressed as mouse chimeric antibodies, with the human ACPA Fab on the mouse IgG2a Fc. Control groups included: naïve wild type (WT), no antibody treatment CAIA (No Ab), and mouse IgG2a isotype control antibody-treated CAIA (IC). ACPAs were first tested in a sub-optimal CAIA model, to evaluate if they could enhance mild joint inflammation in mice, as it has previously been reported^18, 19^. The amount of anti-CII antibody cocktail required for sub-optimal CAIA induction in C57BL/6 mice was optimized in-house and defined as 2.5 mg/mouse, half of the recommended dose for this strain (Suppl. Fig1). Antibodies for each treatment group were injected on day 7 and day 10 post-CAIA induction, and animals were sacrificed on day 14 (Fig. 2A). Clinical scores and paw thickness were assessed every other day to determine arthritis severity (Fig. 2B). To our surprise, both the clinical scores of hind paw inflammation and hind paw thickness were reduced in a statistically significant manner in all ACPA treated groups when compared to controls (Fig. 2B-C). Amelioration of arthritis was seen as early as 2 days post-ACPA antibody injections and it was also evident when comparing the accumulated changes in paw thickness post-antibody injections (Fig. 2C). All ACPA groups presented active reductions of their paw thickness (negative %change) while all CAIA controls had active enlargement of their paws (positive %change). We repeated all experiments by inducing CAIA with the optimal dose of anti-CII antibody cocktail for C57BL/6 (5 mg/mouse) and further refined ACPA specificities to include a representative single mAb for each treatment group (Fig. 2D). The following mAbs were selected: citPAD4+ ACPA: RA64; PAD4-ACPA: RA74; polyspecific ACPA: RA78. A similar ameliorative effect on joint inflammation was seen in all treatment groups when single mAbs were used (Fig. 2E-F). Single ACPAs injected at lower doses (0.5 mg/animal) also ameliorated signs of joint inflammation (Suppl. Fig4). Histological sections of ankle joints showed a marked reduction in signs of arthritis in all ACPA-treated groups (Fig. 3A), with a prominent reduction in joint inflammation (Fig. 3B), cartilage damage (Fig. 3C), and bone erosion (Fig. 3D). Together, our results demonstrate that select, patient-derived ACPA antibodies ameliorate arthritis and reduce joint inflammation in an effector-phase mouse model of arthritis.

**Figure 2.**
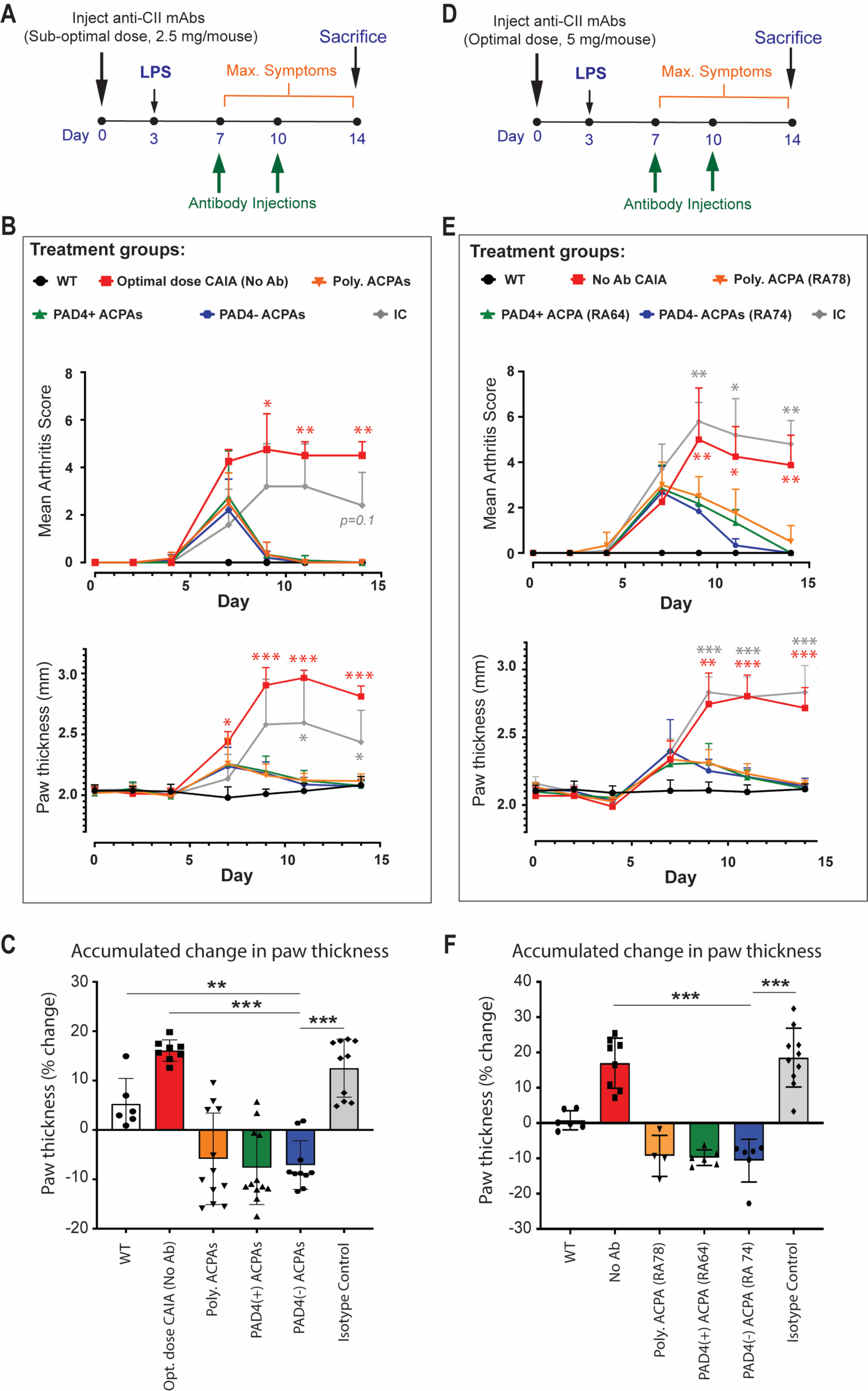
RA monoclonal ACPAs with diverse citrullinated-antigen specificities ameliorate CAIA. **A.** Suboptimal CAIA was induced by injecting C57BL/6 mice with 2.5 mg anti-CII/mouse. Recombinant antibodies were injected on days 7 and 10 post-induction, and animals were sacrificed on day 14. ACPA-treatment groups were formed as follows: (cit-)PAD4+ (RA64, RA65, RA71); PAD4- (RA66, RA74, RA75); and Polyspecific (RA78, RA80, RA82). Control groups: naïve wild-type (WT); CAIA isotype control (IC); Optimal-dose CAIA not injected with recombinant antibodies (NoAb). *n*=3-6 mice/group **B.** The clinical scores of paw inflammation and the hind paw thickness were reduced in all ACPA-treated groups, as early as 2 days post-antibody injection. Compared to isotype controls, mean paw thickness was significantly reduced in all ACPA-treated groups. All ACPA groups were never significantly different from each other and had similar *p* values when compared to controls. By the experiment endpoint, they were not significantly different from WT controls **C.** Accumulated change in hind paw thickness following the first antibody injection. All ACPA-treated groups were significantly different from CAIA controls and showed signs of active amelioration (negative %change). All ACPA groups were not significantly different from each other and had similar *p* values when compared to controls **D.** CAIA was induced by injecting 5 mg anti-CII/animal in C57BL/6 mice. Recombinant antibodies were injected on days 7 and 10 post-induction and animals were sacrificed on day 14. A single ACPA, representative of the combinations tested in ***A***, was injected per group. *n*=3-5 mice/group **E.** The clinical scores of paw inflammation and the hind paw thickness were significantly reduced in all ACPA-treated groups compared to both isotype and no-antibody control animals, and as early as 2 days post-antibody injection. All ACPA groups were never significantly different from each other and had similar *p* values when compared to controls. By the experiment endpoint, they were not significantly different from WT controls **F.** Accumulated change in hind paw thickness following the first antibody injection. All ACPA-treated groups were significantly different from CAIA controls and showed signs of active amelioration (negative %change), suggesting a treatment effect. All ACPA groups were not significantly different from each other and had similar *p* values when compared to controls. *B & E. Only scores for hind legs are shown (1 value/animal, max score: 8). Group mean scores per timepoint +SD are plotted. Both hind paws were included for calculating mean paw thickness (2 values/animal). Group mean paw thickness per timepoint (mm) +SD is plotted. A mixed-effects model with matched values was used for statistical comparisons.* *C & F. Both hind paws were included (2 values/animal). The mean accumulated % change per group +/-SD is plotted. Dots represent individual paws. 1-way ANOVA was used for statistical comparisons.* *B, C, E, F. We only show statistical comparisons for one of the ACPA groups (PAD4-, RA74) vs controls.* *\* = p≤0.05; ** = p≤0.01; *** = p≤0.001*

**Figure 3.**
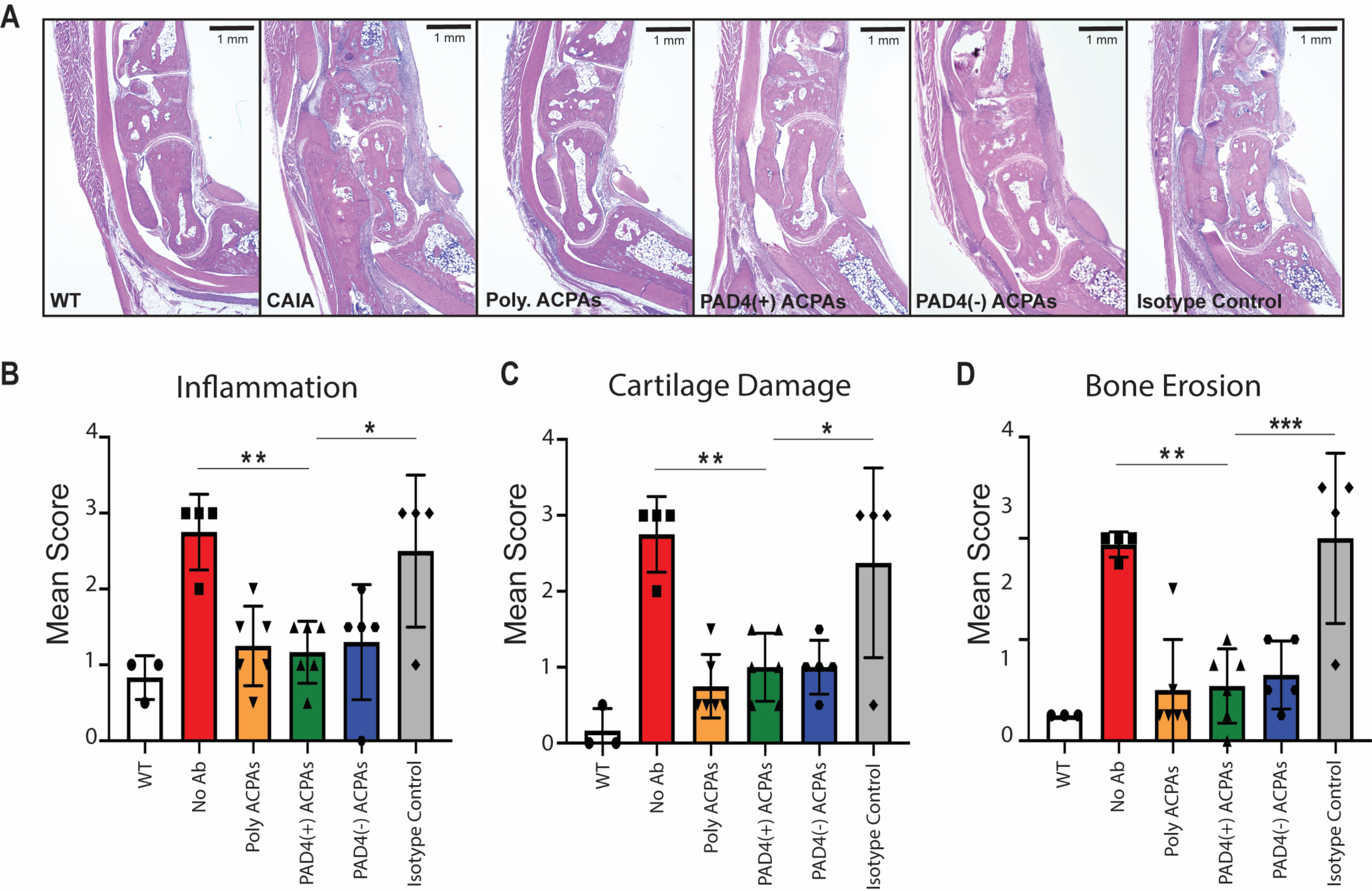
ACPAs reduce histological signs of synovitis and joint damage in CAIA. **A.** Representative histology of ankle joints from ACPA-treated and control groups at endpoint. ACPA groups had less immune-cell infiltration in their joints and better-conserved cartilage and bone structures than isotype and no-antibody (CAIA) controls. Representative joints from wild-type (WT) naïve controls are included for comparison. Tissue sections were stained with hematoxylin-eosin and imaged with a 2× objective lens. Qualitative scoring of **B**) inflammation (0-3), **C**) cartilage damage (0-3), and **D**) bone erosion (0-5). All ACPA-treated groups had significantly lower scores for these parameters than isotype and no-antibody controls (1-Way ANOVA, * = *p*≤0.05; ** = *p*≤0.01; *** = *p*≤0.00*1*). We only show statistical comparisons for one of the ACPA groups (PAD4+ ACPAs) vs controls. Samples were from the experiment summarized in Fig2, A-C. We assessed 1 joint/animal.

### CAIA amelioration by recombinant ACPAs is time dependent

Next, we wanted to determine if the protective properties of ACPAs were dependent on the mouse strain and inflammatory phase of CAIA. To do this we selected two ACPAs from the PAD4 negative group, RA66 and RA74, for further testing in CAIA in DBA/1J mice, a strain that is more prone to develop joint inflammation. First, we tested whether co-injection of ACPAs and the CAIA-inducing antibody cocktail could enhance sub-optimal CAIA, as previously reported^19^. Sub-optimal CAIA was induced by injecting 1 mg anti-CII per mouse in DBA/1J mice, a dose that was optimized in-house (Suppl. Fig1). Recombinant mAbs (1.5 mg/mouse) were injected on day 0, alongside the anti-CII antibody cocktail (Fig. 4A). Co-injection of RA66 or RA74 almost completely abolished development of joint inflammation, as depicted in timelines of clinical scores (Fig. 4B) and paw thickness (Fig. 4C). ACPA-injected groups did not show any clinical signs of CAIA throughout the experiment and were statistically different from both isotype control and no-antibody CAIA animals from day 11 onwards (*p*<0.05). The total accumulated change in paw thickness clearly illustrates how CAIA was prevented in ACPA-treated groups, which were statistically different from CAIA controls and did not actively increase their paw size (Fig. 4D). Thus, co-injection of select monoclonal ACPAs protected mice from developing CAIA arthritis.

**Figure 4.**
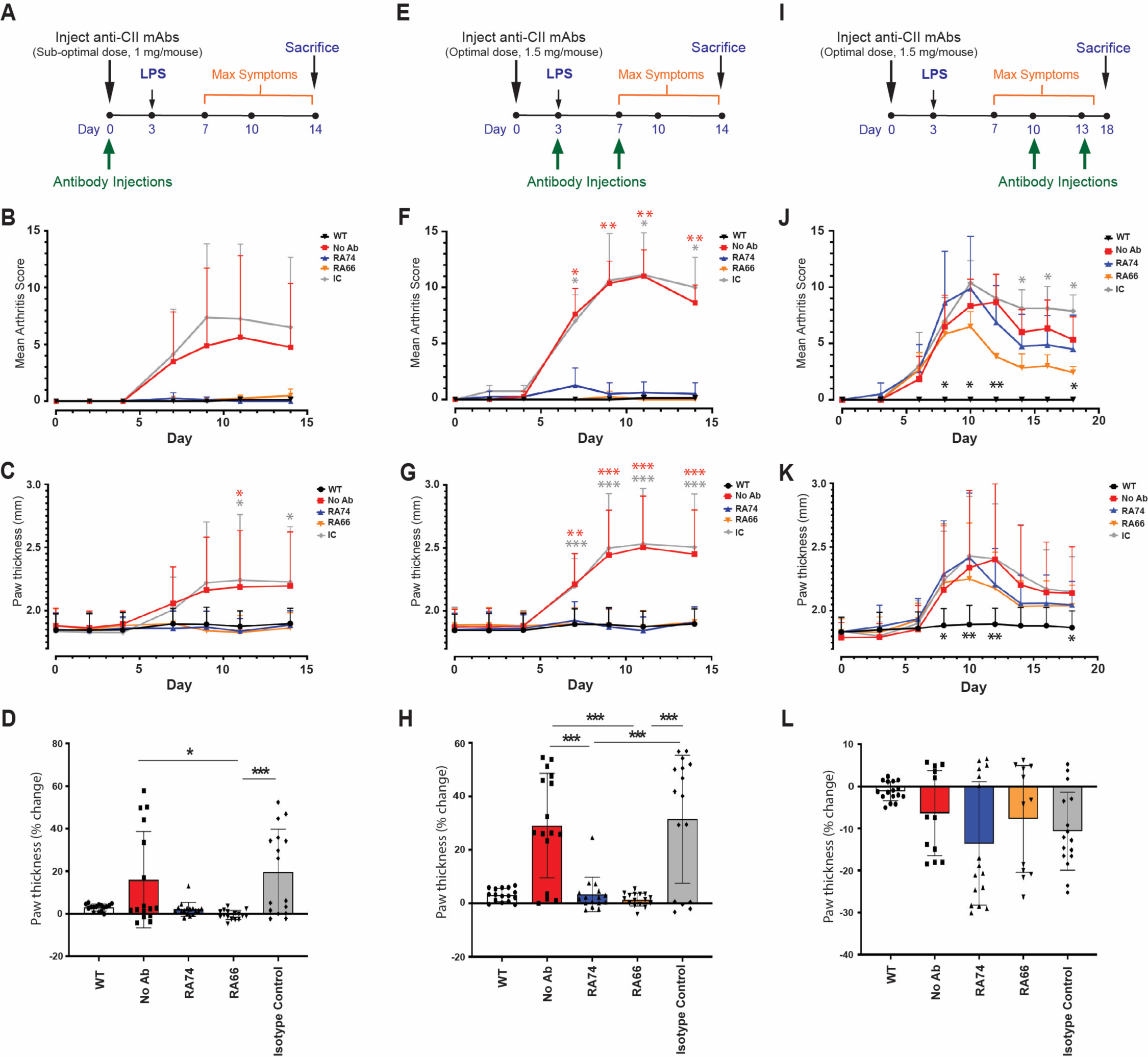
ACPAs’ amelioration of CAIA is time dependent. **A.** Co-injection of ACPAs and anti-CII cocktail. Sub-optimal CAIA was induced by injecting 1 mg anti-CII/mouse in DBA/1J mice. Recombinant antibodies were injected on day 0, alongside the anti-CII antibody cocktail, and animals were sacrificed on day 14. Two recombinant ACPAs were tested in this and subsequent CAIA experiments: RA66 and RA74 (both PAD4-ACPAs). Control groups included: naïve wild-type (WT), CAIA isotype control (IC), and CAIA not injected with recombinant antibodies (No Ab). *n*=4 mice/group. Co-injection of ACPAs completely abolished the development of joint inflammation, as depicted in the clinical scores (**B**) and paw thickness (**C**) timelines. ACPA-injected groups did not show any clinical signs of CAIA throughout the experiment. Paw thickness was significantly reduced for RA66-injected animals, compared to both isotype control and no-antibody CAIA groups, from day 10 onwards. Repeated measures 2-way ANOVA was used for statistical analysis **D**. Accumulated change in paw thickness following the first antibody injection. ACPA-treated groups were significantly different from CAIA controls and showed no accumulated paw inflammation. They were not significantly different from WT controls. Kruskal-Wallis tests were used for comparisons **E.** ACPAs in early-phase CAIA. CAIA was induced by injecting 1.5 mg anti-CII/mouse in DBA/1J mice. Recombinant antibodies were injected on days 3 and 7 post-induction and animals were sacrificed on day 14. n=4 mice/group. As shown in the clinical scores (**F**) and paw thickness (**G**) timelines, ACPAs fully prevented the development of joint inflammation when injected at early stages of CAIA. ACPA-treated groups had significantly lower clinical scores and paw thickness than both isotype and no-antibody control animals from day 7 onwards, and they were always undistinguishable from WT controls. Repeated measures 2-way ANOVA was used for statistical analysis **H**. Accumulated change in paw thickness following the first antibody injection. All ACPA-treated groups were significantly different from CAIA controls and showed no signs of paw inflammation. They were not significantly different from WT controls. 1-way ANOVA was used for statistical comparisons **I**. ACPAs in late-stage CAIA. CAIA was induced by injecting 1.5 mg anti-CII/mouse in DBA/1J mice. Recombinant antibodies were injected on days 10 and 13 post-induction and animals were sacrificed on day 18. n=4 mice/group. Clinical scores (**J**) and paw thickness (**K**) were decreased after injection of ACPAs on day 10, compared to no antibody and to isotype controls, though only significantly for the clinical scores of RA66-treated mice. Injection of ACPAs on day 13 did not alter the resolution phase of CAIA in any capacity. ACPA-treated groups were never significantly different from each other, but they remained significantly different from WT controls for most of the experiment. A mixed-effects model with matched values was used for statistical comparisons **L**. Accumulated change in paw thickness following the first antibody injection. All CAIA groups experienced reductions in their accumulated paw thickness, regardless of treatment, indicating a recovery phase for CAIA. ACPA-treated groups were not significantly different from CAIA controls. Kruskal-Wallis tests were used for comparisons. *B,F,J All paws were included in the calculation of mean clinical scores (1 value/animal, max score: 16). Group mean scores per timepoint +SD are plotted.* *C,G,K All paws were included for calculating mean paw thickness (4 values/animal). Group mean paw thickness per timepoint (mm) +SD is plotted.* *D,H,L All paw were included for calculating accumulated % change in paw thickness (4 values/animal). The mean accumulated % change per group +/-SD is plotted. Dots represent individual paws.* *B, C, D, F, G, H, J, K, L. We only show statistical comparisons for one of the ACPA groups (RA74) vs controls. * = p≤0.05; ** = p≤0.01; *** = p≤0.001*

Subsequently, we tested if ACPAs could prevent optimal-dose CAIA when injected in early-phases of the model. We induced CAIA with 1.5 mg anti-CII/mouse on day 0 and mAbs were injected on day 3 and day 7 post-CAIA induction (Fig. 4E). Timelines of clinical scores and paw thickness showed that delayed ACPA injections during the early phase of CAIA fully prevented the development of joint inflammation (Fig. 4F-G). ACPA-treated groups had significantly lower clinical scores and paw thickness than both isotype (p<0.05 and p<0.001, respectively) and no-antibody control animals (p<0.01 and p<0.001, respectively) from day 7 onwards, and were undistinguishable from WT controls throughout the experiment. The accumulated changes in paw thickness further emphasize that ACPAs protected animals from developing joint inflammation when injected on day 3 and day 7 after CAIA induction (Fig. 4H). All ACPA-treated groups were significantly different from CAIA controls and showed no signs of paw inflammation (p<0.001). These findings show that early phase injections of select monoclonal ACPAs were able to protect animals from developing optimal-dose CAIA.

Next, we determined if ACPAs can lessen the effects of CAIA at a later stage of disease development. We repeated the above experiments, but delayed ACPA injections to day 10 and day 13 post-CAIA induction and sacrificed animals on day 18 (Fig. 4I). Clinical scores (Fig. 4J) and paw thickness (Fig. 4K) were lower after ACPA injections on day 10, compared to no antibody and isotype controls, though only significantly for RA66. Injection of ACPAs on day 13 did not seem to alter the late phase of CAIA in any capacity. In addition, late-phase injections of ACPAs did not produce statistically significant differences in the accumulated changes in paw thickness post-antibody-injection, all groups presented reductions consistent with a resolution phase (Fig. 4L). Altogether, these results indicate that, while select ACPAs are effective at both preventing the onset and reducing the severity of CAIA when injected early in the model, they do not drastically influence the resolution phase of the model.

## DISCUSSION

Anti-citrullinated protein antibodies (ACPAs), along with rheumatoid factor, are hallmark autoantibodies in rheumatoid arthritis (RA). They are present in at-risk individuals years prior the clinical diagnosis of RA^13^ and are associated with severe bone damage in patients with established RA^11^. While several studies support a pathogenic role for ACPAs in RA, they are predominantly *in vitro* studies^32^. Publications addressing the direct role of ACPAs in animal models of joint inflammation are rare and have delivered mixed results. For instance, injecting mice with anti-citVimentin ACPAs triggered osteoclastogenesis and osteopenia^17, 33^ or induced pain-like behavior^16^, but did not produce significant joint inflammation. Most studies describing a pathogenic role for ACPAs in animal models used non-patient derived recombinant antibodies and reported enhancement of a pre-existing, mild joint inflammation^18, 19, 34^. In this study, we tested a combination of 9 recombinant, patient-derived ACPAs with diverse citrullinated antigen (citAgs) specificities in collagen antibody-induced arthritis (CAIA), an acute-phase murine model of joint inflammation that engages primarily the innate immune system and relies heavily on neutrophil activation^35^. We showed that injection of recombinant ACPAs at early stages of CAIA both prevented and ameliorated paw inflammation, irrespective of the citAg specificities involved. ACPAs, however, did not affect the normal course of CAIA when injected at later stages of the model. Of note, CAIA was ameliorated despite using ACPAs grafted in an inflammatory mouse IgG2a Fc.

The present study highlights the time-dependency of the ACPA-mediated amelioration of CAIA (Suppl. Fig5). Kuhn *et al.* reported that co-injection of recombinant anti-citrullinated fibrinogen ACPAs in a sub-optimal model of CAIA enhanced arthritis symptoms^19^. We replicated these experimental conditions with select ACPAs derived from RA patients but encountered full prevention of CAIA instead. Kuhn’s antibodies were mouse CIA hybridoma-derived and included one IgM and two IgGs, selected for their binding to citrullinated fibrinogen. The ACPAs tested here were all grafted onto mouse IgG2a Fc and included reactivity towards citrullinated fibrinogen as well as to other citAgs. One possible explanation for these discrepant outcomes may be the ACPAs’ diverse citrullinated antigen specificities, which might have triggered anti-inflammatory mechanisms that overcame the pro-inflammatory effect of anti-citrullinated fibrinogen reactivities. In addition, we tested patient-derived ACPAs with high affinity for citrullinated targets, as opposed to mouse-derived ACPAs with undisclosed affinities. Furthermore, our results highlight the strength of ACPAs’ preventive effect in CAIA; a single ACPA injection given in conjunction with the CAIA-induction cocktail fully abrogated the development of arthritis. This is further supported by the complete prevention of joint inflammation when ACPAs were injected on days 3 and 7 of CAIA. Interestingly, late-stage CAIA was not affected by recombinant ACPAs, suggesting that citAgs, or the cells mediating ACPAs’ ameliorative effect, are not critically involved in the recovery phase of CAIA.

To further define the role of citAg specificities in the ACPA-mediated amelioration of CAIA, we used homogeneous antibody isotypes (mouse IgG2a), produced under standard conditions and tested at equal concentrations. They differed only in their binding specificities to certain citAgs, which was the rationale for combining them into test groups. As illustrated by this study, all ACPA combinations ameliorated early CAIA, providing evidence against a fundamental role for citAgs specificities in the anti-inflammatory effects of these ACPAs. The antigen targets of this set of recombinant ACPA mAbs included: citPAD4, histones (particularly H3 and H4, both native and citrullinated), and citFibrinogen. Binding to PAD4 was more restricted and presented a more “private” type of recognition, primarily in one of the clonal families. Anti-PAD4 antibodies, especially those that cross-react to PAD3, have been linked to worse disease outcomes in RA^30^, but we did not observe enhanced inflammation in CAIA when injecting ACPAs that included PAD4/3 specificities, on the contrary, they ameliorated CAIA. These contradictory findings might be explained by additional citAgs specificities included in the selected ACPAs but, overall, they opposed the notion of a strong pro-inflammatory effect for anti-PAD4 antibodies. In contrast to PADs, binding to histones was observed in all clonal families studied here. Histones are prominent citAgs in neutrophil extracellular traps (NETs)^36^, thus, one possible explanation for the generalized ameliorative effect of this ACPA cohort is through interactions with NETs, a major source of citrullinated histones in inflamed synovium^37^. Indeed, representative ACPAs from our cohort bound to *in vitro*-generated NETs and co-localized with citHistones (data not shown), which has also been shown for other ACPAs ^38^.

NETs were initially described in 2004^39^, and have been well-established to play a pivotal role in controlling microbial infections. However, it has also become evident that chronic, dysregulated exposure to NETs is detrimental in several inflammatory conditions, including autoimmune diseases such as SLE, ANCA vasculitis, and RA^40, 41^. Chirivi *et al.* showed that ACPA injections ameliorated inflammation in several animal models in which NETs are implicated, including CAIA^22^. The ACPAs’ effect was Fc-dependent and attributed to both inhibition of NET formation and enhanced macrophage NET clearance. ACPAs from Chirivi’s study were mouse-zised versions (mouse IgG2a Fc grafted onto human Fab) of library-derived human anti-citH4 antibodies. Recently, Raposo *et al.* described similar effects on early CAIA animals treated with patient-derived ACPAs^20^. This effect was observed for some of the tested ACPAs and was not associated with their fine specificities, though binding to full protein histones and PADs were not reported. In addition, He at al. showed that CAIA was prevented by co-injection of an ACPA targeting citrullinated alpha enolase (citENO1)^21^. This effect was mediated by the interaction between immune complexes of ACPA-citENO1 and FCGR2B in macrophages, which promoted IL-10 secretion and reduced osteoclastogenesis. Our study corroborates and expands these findings with affinity-matured, patient-derived ACPAs that bind citrullinated histones and additional citrullinated antigen specificities, including citrullinated PADs. Importantly, we highlight the preventive, rather than therapeutic, nature of ACPAs’ effects in CAIA.

The limitations of this study include testing 9 different ACPAs that were all derived from a single RA patient. Thus, generalizations of ACPAs’ anti-inflammatory effects would require expanding these studies with mAbs from additional patients or absorbed ACPAs from RA sera. In addition, the predominantly polyspecific nature of the ACPAs here studied prevents a proper parsing of the different citAg-specificities’ contributions to the amelioration of CAIA. Conducting systematic experiments with single-antigen specific ACPAs might advance our understanding of the relative relevance of different citAgs, and their respective autoantibodies, in arthritis.

The deposition of immune complexes of ACPAs and citAgs in affected joints is largely considered as pro-inflammatory^42^. This is, however, a generalization as the nature of immune complexes and their effects *in vivo* are heterogeneous and rely on various factors, such as antigen type(s), relative concentrations, valences, the antibody isotypes involved, Fc glycosylation levels, interactions with Fc receptors, etc^43–45^. In addition, the formation of immune complexes is fundamental for the clearance of target antigens by the innate immune system^46^. Depending on the antigens involved, this immune-complex mediated clearance could contribute to dampen inflammation.

The most prevalent hypothesis for ACPAs’ mechanism of action in RA is that their binding to citAgs (e.g. in NETs) generate immune-complexes that accumulate in the synovium and activate infiltrating immune cells, thereby promoting inflammation. An alternative hypothesis is that this ACPA-mediated immune activation enhances NET clearance, or blockade, thereby reducing inflammation. Definitive evidence for or against these potential mechanisms is lacking. Although this study does not address molecular mechanisms, the ACPAs’ anti-inflammatory effects reported herein favor a NETs/citAgs neutralization hypothesis. Given their load of inflammatory mediators, NETs are an important source of damage-associated molecular-patterns (DAMP) molecules, such as histone H3, cell-free DNA, HMGB1, and S100 proteins^47^. Importantly, because of PAD activation and extracellular exposure during NETosis, many NET-associated DAMPs are citrullinated, which renders them even more pro-inflammatory than their native forms^48, 49^. In this regard, ACPAs could serve as countermeasures for mitigating NET-mediated inflammation. For instance, it has been shown that citH3 levels are elevated in septic patients^50, 51^, and that injections of recombinant anti-citH3 antibodies improved survival in animal models of septic shock^52, 53^. Thus, blocking citrullinated DAMPs with antibodies can dampen sudden, generalized inflammation. Arguably, a similar process might be occurring with ACPAs in acutely inflamed CAIA joints, though in a more localized fashion. Of note, ACPA-injected animals displayed an improved overall appearance and body weight compared to controls, suggesting that ACPAs might have counteracted the LPS-induced, generalized inflammation of CAIA.

Previous studies reported the potential of germline-reverted ACPA antibodies to conserve their binding to citrullinated epitopes^5, 54^. Indeed, we also observed binding to citPAD4 in germline-reverted ACPAs from one of the clonal families (CF2). Thus, it is possible that the immune system generates “ACPA natural autoantibodies” that subsequently undergo T cell-promoted affinity maturation. In line with this, it has been reported that patient-derived, pentameric IgM ACPAs bind to a variety of post-translationally modified self-antigens, including citAgs, and that these properties are conserved within their germline ancestry^55^. Transgenic mice expressing IgM natural antibodies^56^, or mice injected with anti-dsDNA IgM antibodies^57^, were protected from nephritic damage in spontaneous models of lupus. Further, certain ACPAs cross-react with citrullinated bacterial antigens^58^ and germline-reverted cross-reactive ACPAs maintain their specificities for bacteria but not for citAgs^59^, suggesting that there is a complex interplay between anti-microbial responses and the reactivity towards post-translationally modified self-proteins. Of note, most of the ACPAs characterized here were derived from IgA plasmablasts^5^, implying a possible link between mucosal immunity and the development of protective ACPAs.

In summary, we show that patient-derived, high-affinity ACPAs, polyspecific for several RA-associated citAgs including citPAD4, prevented and/or ameliorated the acute phase of CAIA. The same recombinant ACPAs did not affect the severity of CAIA at late stages. Taking into account that ACPAs are present years prior to the onset of clinical RA, and the ACPA-mediated amelioration of joint inflammation reported herein, we speculate that ACPAs might play a protective role in reducing joint inflammation in pre-clinical RA. In established RA, epitope spreading, isotype switching, and the altered immune cell profile of the inflamed synovium may render protective ACPAs ineffective or pathogenic. Discerning the mechanisms underlying the ACPA-mediated amelioration of CAIA, as well as the function of ACPAs in pre-RA and established RA, could fundamentally reshape our understanding of the role that ACPAs play in RA.

## ACKNOWLEDGEMENTS

We would like to extend our recognition to Benjamin Franco, for his valuable help in all animal experiments, to Nithya Lingampalli, for her contributions in running planar antigen arrays, and to the following Gilead Sciences Inc. researchers: Magdeleine Hung, Sabrina Lu, Elizabeth Chen, Ian Scott, Debi Jin, and Upasana Mehra for their help with the production of recombinant ACPAs and initial characterization. This study was partially supported by funding from Gilead Sciences Inc.

---

**Supplemental Figure 1.**
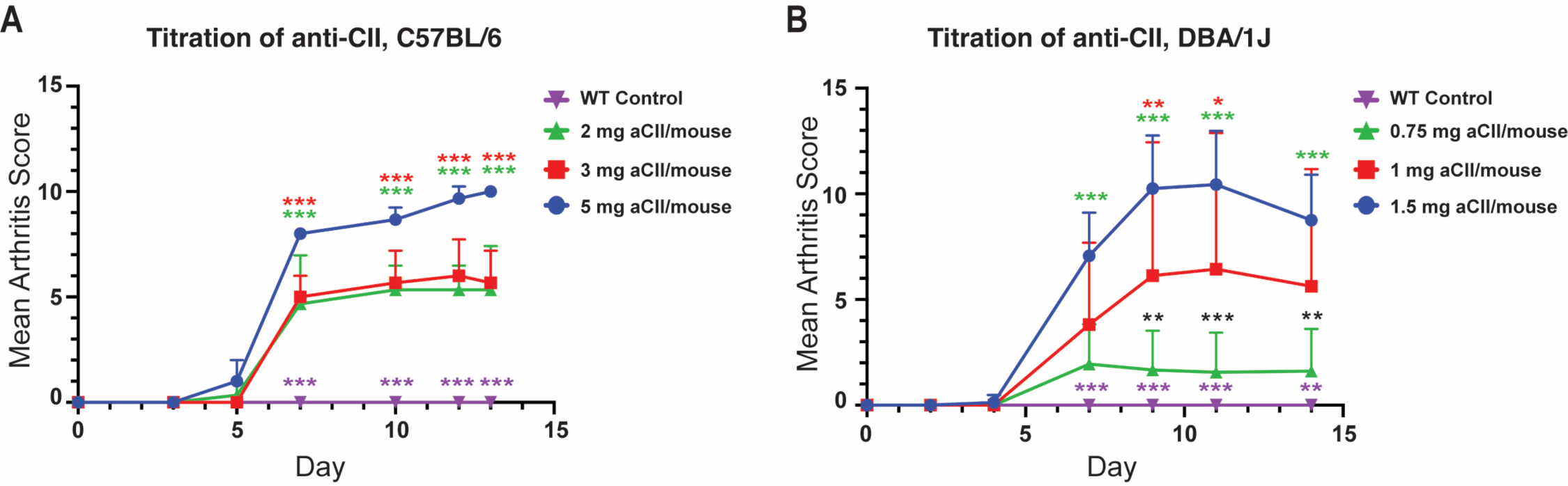
Titration of anti-CII cocktail in C57BL/6 and DBA/1J mice. **A.** C57BL/6 mice were injected with different doses of anti-CII/mouse: 5 mg (recommended dose), 3 mg, and 2 mg. **B.** DBA/1J mice were injected with different doses of anti-CII/mouse: 1.5 mg (recommended dose), 1 mg, and 0.75 mg. Naïve wild-type (WT) controls were not injected with anti-CII antibodies. Timelines depict mean arthritis scores throughout the experiment (maximum score:16). Animals were sacrificed on day 14. In **A**, all doses were significantly different from WT controls from day 7 onwards and there were no significant differences between sub-optimal doses throughout the experiment (purple asterisks). In **B**, the 1 mg aCII/mouse group was significantly different from WT controls from day 9 onwards, while the 0.75 mg aCII/mouse group was never significantly different from WT controls (purple asterisks). 2-way ANOVAs were used for statistical analysis, colored asterisks indicate significant differences between sub-optimal doses and the corresponding optimal dose for the strain, while black asterisks represent significant differences between sub-optimal doses. *n*=3-5 mice/group.

**Supplemental Figure 2.**
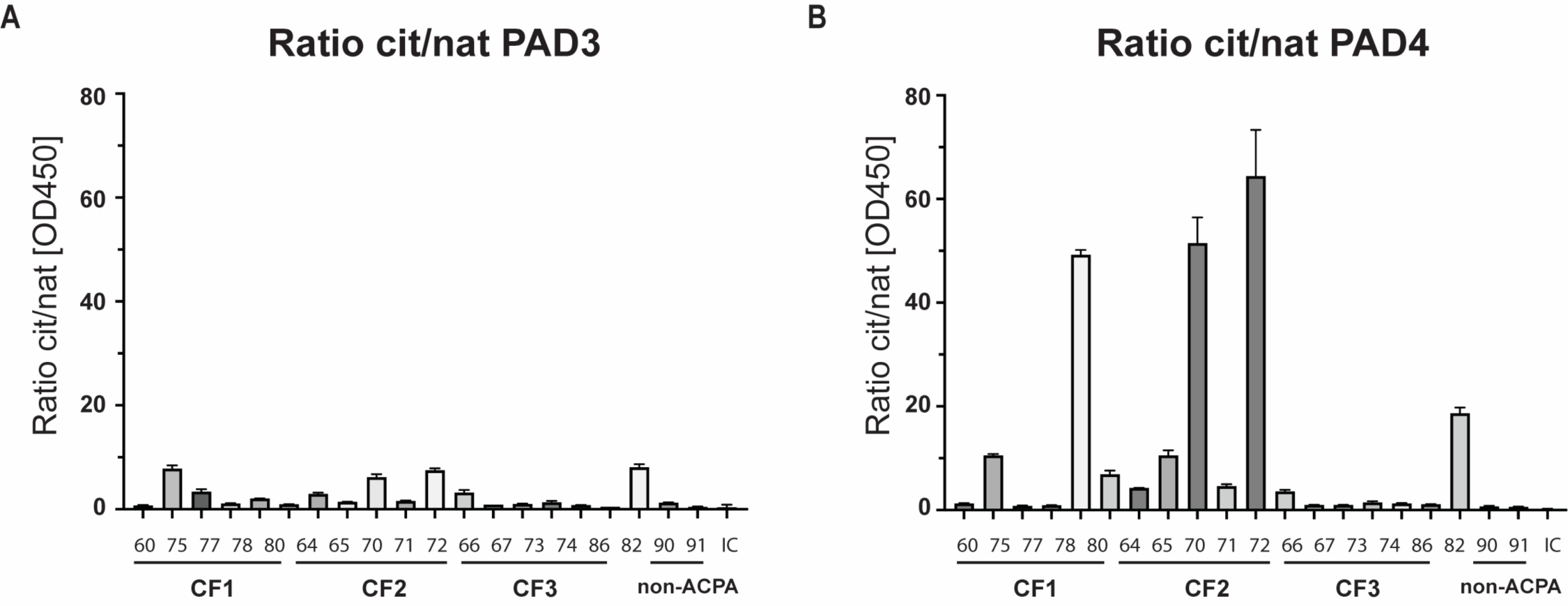
Ratios for the binding of ACPAs to citrullinated/native PAD3 and PAD4. Recombinant native and auto-citrullinated PAD3 and PAD4 were coated on ELISA plates and incubated with monoclonal antibodies (mAbs). The ratios of blank-corrected ODs for citrullinated/native PAD3 **(A)** and PAD4 **(B)** are plotted. Both recombinant PADs were from Cayman Chemical. Auto-citrullinated PADs were generated by incubating PAD enzymes in the presence of 2 mM calcium chloride and 2 mM DTT for 2 hours at 37C. Antigens were coated at 2 μg/mL, in carbonate buffer, and antibodies were tested at 15 μg/mL. Each condition was tested in duplicate. RA90 and RA91 are patient-derived non-ACPA mAb controls. *IC: isotype control antibody; cit: citrullinated; nat: native*

**Supplemental Figure 3.**
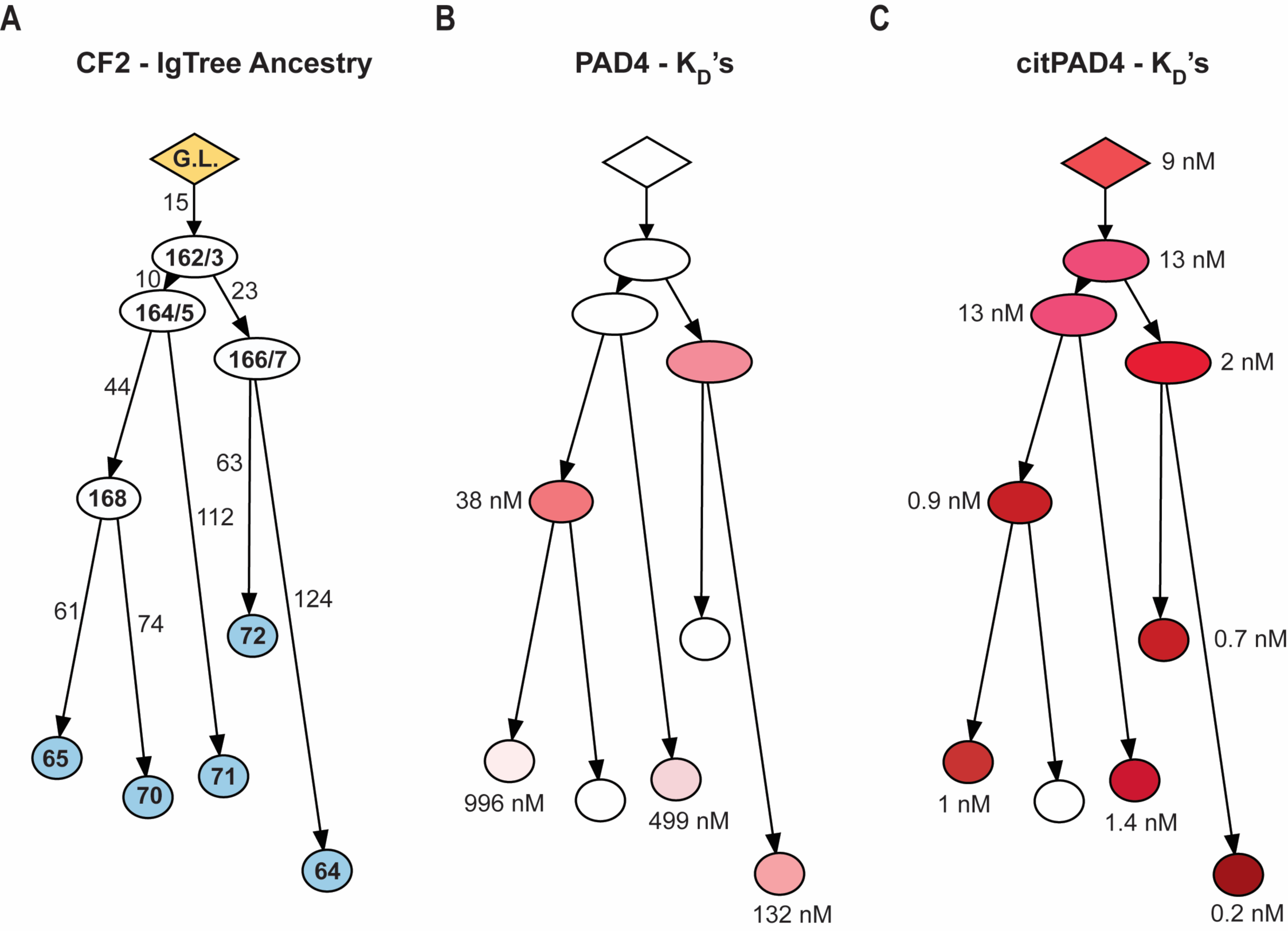
K_D_’s of germline-reverted ACPAs for their binding to PAD4 and citPAD4. Equilibrium dissociation constants (K_D_’s) were determined by bio-layer interferometry (Octet system, ForteBio/Sartorius) for plasmablast-sequenced, affinity-matured ACPAs from one clonal family (CF2) and their ancestral and germline-reverted antibodies **A.** The immunoglobulin phylogenetic tree for CF2 and its ancestral members was generated by IgTree^31^, numbers in the branches indicate number of mutations between members. K_D_’s were calculated for the binding of ancestral antibodies to native **(B)** and auto-citrullinated **(C)** recombinant PAD4 and are reported next to their corresponding antibody circle. Color intensities are proportional to binding affinities (higher affinity: darker color). Antibody circles left blank had no binding. K_D_’s are reported in nM and were computed by globally fitting at least 2 serially diluted ligand curves per kinetic assay. Recombinant PAD4 was from Cayman Chemical. Auto-citrullinated PAD4 were generated by incubating the enzyme in the presence of 2 mM calcium chloride and 2 mM DTT for 2 hours at 37C. *Yellow: germline sequence (G.L.); White: ancestral/parent antibodies; Blue: affinity-matured antibodies; CF: clonal family*

**Supplemental Figure 4.**
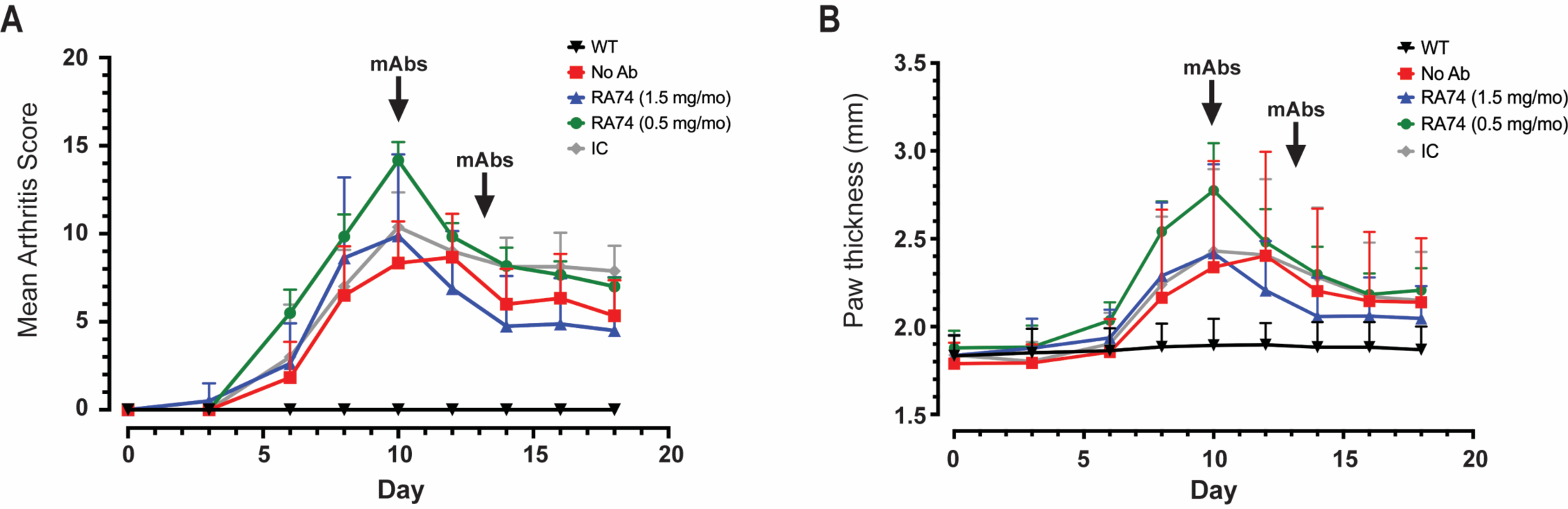
ACPAs ameliorate CAIA at multiple doses. CAIA was induced in DBA/1J mice by injecting 1.5 mg anti-CII cocktail/mouse. Recombinant ACPA (RA74) and isotype control (IC) antibodies were injected on days 10 and 13. For RA74, two doses were tested: 1.5 mg/mouse/injection and 0.5 mg/mouse/injection. The isotype control antibody was injected at 1.5 mg/mouse/injection. No antibody-treated CAIA controls (No Ab) were injected with vehicle, while wild-type (WT) naïve controls did not receive any antibody injection. Timelines of mean clinical scores **(A)** and mean paw thickness **(B)** are shown. After the first antibody injection (day 10), there were no significant differences between the two ACPA doses tested, for both parameters here displayed. Of note, both ACPA groups (0.5 and 1.5 mg/mouse) showed sharper reductions of their clinical scores and paw thickness after the first antibody injection (day 10) compared to CAIA controls, though not significantly. No differences were observed after the second antibody injection (day 13). 2-way ANOVAs were used for statistical analysis. *n*=4 mice/group.

**Supplemental Figure 5.**
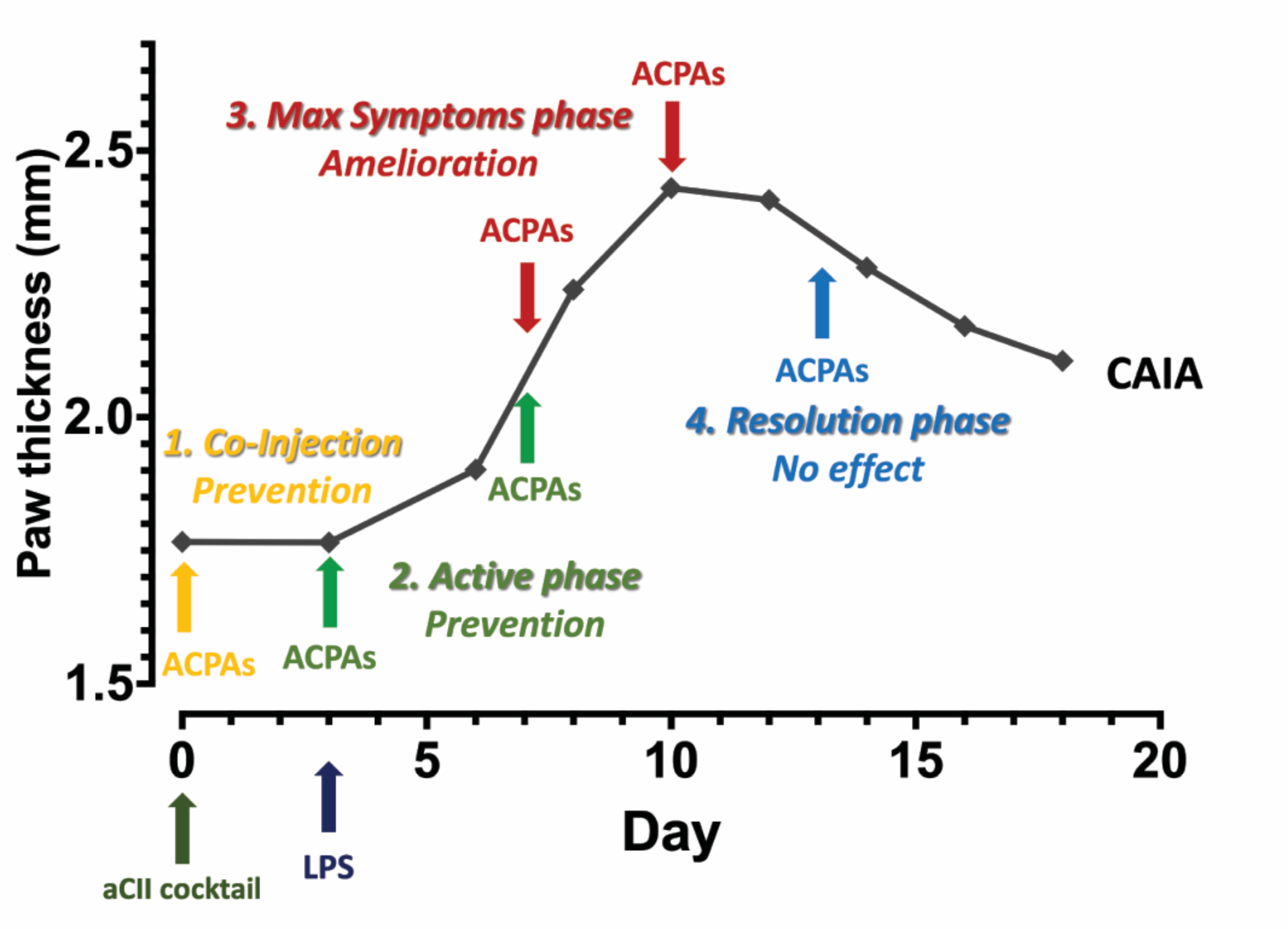
Summary of ACPAs’ effects at different stages of CAIA.

1. *Co-injection of ACPAs and anti-Collagen type II antibodies (aCII cocktail).* Injection of ACPAs along with the collagen-antibody induced arthritis (CAIA) induction cocktail (day 0) led to full prevention of CAIA.
2. *ACPAs’ injection at early stages of CAIA.* When delivered on day 3 (along with the LPS injections) and day 7, ACPAs also prevented the development of CAIA.
3. *ACPAs’ injection at the maximum-symptoms phase of CAIA.* Injection of ACPAs after the onset of arthritis (day 7 and day 10) significantly ameliorated joint inflammation but did not result in full recovery from CAIA.
4. *ACPAs’ injection at the resolution phase of CAIA.* ACPAs injected during the late stages of CAIA (day 13) did not significantly alter the normal resolution of joint inflammation observed in this phase.

## Notes

### Competing Interest Statement

- GMO and ANM are employed by Gilead Sciences, Inc.
- WHR is a consultant to, owns equity in, and serves on the Board of Directors of Atreca, Inc.
- This study was partially supported by funding from Gilead Sciences, Inc.

## REFERENCES

1. Ronnelid, J., Turesson, C. & Kastbom, A. Autoantibodies in Rheumatoid Arthritis - Laboratory and Clinical Perspectives. Front Immunol 12, 685312, doi:10.3389/fimmu.2021.685312 (2021).

2. Ciesielski, O. et al. Citrullination in the pathology of inflammatory and autoimmune disorders: recent advances and future perspectives. Cell Mol Life Sci 79, 94, doi:10.1007/s00018-022-04126-3 (2022).

3. Catrina, A., Krishnamurthy, A. & Rethi, B. Current view on the pathogenic role of anti-citrullinated protein antibodies in rheumatoid arthritis. RMD Open 7, doi:10.1136/rmdopen-2020-001228 (2021).

4. Ge, C. et al. Structural Basis of Cross-Reactivity of Anti-Citrullinated Protein Antibodies. Arthritis Rheumatol 71, 210–221, doi:10.1002/art.40698 (2019).

5. Kongpachith, S. et al. Affinity Maturation of the Anti-Citrullinated Protein Antibody Paratope Drives Epitope Spreading and Polyreactivity in Rheumatoid Arthritis. Arthritis Rheumatol 71, 507–517, doi:10.1002/art.40760 (2019).

6. Lee, D. M. & Schur, P. H. Clinical utility of the anti-CCP assay in patients with rheumatic diseases. Ann Rheum Dis 62, 870-874, doi:10.1136/ard.62.9.870 (2003).

7. Payet, J. et al. ACPA-positive primary Sjogren’s syndrome: true primary or rheumatoid arthritis-associated Sjogren’s syndrome? RMD Open 1, e000066, doi:10.1136/rmdopen-2015-000066 (2015).

8. Ahn, S. S., Pyo, J. Y., Song, J. J., Park, Y. B. & Lee, S. W. Anti-Citrullinated Peptide Antibody Expression and Its Association with Clinical Features and Outcomes in Patients with Antineutrophil Cytoplasmic Antibody-Associated Vasculitis. Medicina (Kaunas) 58, doi:10.3390/medicina58040558 (2022).

9. van Zanten, A. et al. Presence of anticitrullinated protein antibodies in a large population-based cohort from the Netherlands. Ann Rheum Dis 76, 1184–1190, doi:10.1136/annrheumdis-2016-209991 (2017).

10. Nienhuis, R. L. & Mandema, E. A NEW SERUM FACTOR IN PATIENTS WITH RHEUMATOID ARTHRITIS; THE ANTIPERINUCLEAR FACTOR. Ann Rheum Dis 23, 302–305, doi:10.1136/ard.23.4.302 (1964).

11. Syversen, S. W. et al. Prediction of radiographic progression in rheumatoid arthritis and the role of antibodies against mutated citrullinated vimentin: results from a 10-year prospective study. Ann Rheum Dis 69, 345–351, doi:10.1136/ard.2009.113092 (2010).

12. Conforti, A. et al. Beyond the joints, the extra-articular manifestations in rheumatoid arthritis. Autoimmun Rev 20, 102735, doi:10.1016/j.autrev.2020.102735 (2021).

13. Sokolove, J. et al. Autoantibody epitope spreading in the pre-clinical phase predicts progression to rheumatoid arthritis. PLoS One 7, e35296, doi:10.1371/journal.pone.0035296 (2012).

14. Nielen, M. M. et al. Specific autoantibodies precede the symptoms of rheumatoid arthritis: a study of serial measurements in blood donors. Arthritis Rheum 50, 380–386, doi:10.1002/art.20018 (2004).

15. Serdaroglu, M., Cakirbay, H., Deger, O., Cengiz, S. & Kul, S. The association of anti-CCP antibodies with disease activity in rheumatoid arthritis. Rheumatol Int 28, 965–970, doi:10.1007/s00296-008-0570-3 (2008).

16. Wigerblad, G. et al. Autoantibodies to citrullinated proteins induce joint pain independent of inflammation via a chemokine-dependent mechanism. Ann Rheum Dis 75, 730–738, doi:10.1136/annrheumdis-2015-208094 (2016).

17. Harre, U. et al. Induction of osteoclastogenesis and bone loss by human autoantibodies against citrullinated vimentin. J Clin Invest 122, 1791–1802, doi:10.1172/JCI60975 (2012).

18. Sohn, D. H. et al. Local Joint inflammation and histone citrullination in a murine model of the transition from preclinical autoimmunity to inflammatory arthritis. Arthritis Rheumatol 67, 2877–2887, doi:10.1002/art.39283 (2015).

19. Kuhn, K. A. et al. Antibodies against citrullinated proteins enhance tissue injury in experimental autoimmune arthritis. J Clin Invest 116, 961–973, doi:10.1172/JCI25422 (2006).

20. Raposo, B. et al. Divergent and dominant anti-inflammatory effects of patient-derived anticitrullinated protein antibodies (ACPA) in arthritis development. Ann Rheum Dis, doi:10.1136/ard-2022-223417 (2023).

21. He, Y. et al. A subset of antibodies targeting citrullinated proteins confers protection from rheumatoid arthritis. Nat Commun 14, 691, doi:10.1038/s41467-023-36257-x (2023).

22. Chirivi, R. G. S. et al. Therapeutic ACPA inhibits NET formation: a potential therapy for neutrophil-mediated inflammatory diseases. Cell Mol Immunol 18, 1528–1544, doi:10.1038/s41423-020-0381-3 (2021).

23. Nair, N. et al. VP4- and VP7-specific antibodies mediate heterotypic immunity to rotavirus in humans. Sci Transl Med 9, doi:10.1126/scitranslmed.aam5434 (2017).

24. Tan, Y. C. et al. High-throughput sequencing of natively paired antibody chains provides evidence for original antigenic sin shaping the antibody response to influenza vaccination. Clin Immunol 151, 55–65, doi:10.1016/j.clim.2013.12.008 (2014).

25. Robinson, W. H. et al. Autoantigen microarrays for multiplex characterization of autoantibody responses. Nat Med 8, 295-301, doi:10.1038/nm0302-295 (2002).

26. Hueber, W. et al. Antigen microarray profiling of autoantibodies in rheumatoid arthritis. Arthritis Rheum 52, 2645–2655, doi:10.1002/art.21269 (2005).

27. Cunningham, O., Scott, M., Zhou, Z. S. & Finlay, W. J. J. Polyreactivity and polyspecificity in therapeutic antibody development: risk factors for failure in preclinical and clinical development campaigns. MAbs 13, 1999195, doi:10.1080/19420862.2021.1999195 (2021).

28. Hayer, S. et al. ’SMASH’ recommendations for standardised microscopic arthritis scoring of histological sections from inflammatory arthritis animal models. Ann Rheum Dis 80, 714–726, doi:10.1136/annrheumdis-2020-219247 (2021).

29. Thiam, H. R. et al. NETosis proceeds by cytoskeleton and endomembrane disassembly and PAD4-mediated chromatin decondensation and nuclear envelope rupture. Proc Natl Acad Sci U S A 117, 7326–7337, doi:10.1073/pnas.1909546117 (2020).

30. Darrah, E. et al. Erosive rheumatoid arthritis is associated with antibodies that activate PAD4 by increasing calcium sensitivity. Sci Transl Med 5, 186ra165, doi:10.1126/scitranslmed.3005370 (2013).

31. Barak, M., Zuckerman, N. S., Edelman, H., Unger, R. & Mehr, R. IgTree: creating Immunoglobulin variable region gene lineage trees. J Immunol Methods 338, 67–74, doi:10.1016/j.jim.2008.06.006 (2008).

32. Toes, R. & Pisetsky, D. S. Pathogenic effector functions of ACPA: Where do we stand? Ann Rheum Dis 78, 716–721, doi:10.1136/annrheumdis-2019-215337 (2019).

33. Engdahl, C. et al. Periarticular Bone Loss in Arthritis Is Induced by Autoantibodies Against Citrullinated Vimentin. J Bone Miner Res 32, 1681–1691, doi:10.1002/jbmr.3158 (2017).

34. Ho, P. P. et al. Autoimmunity against fibrinogen mediates inflammatory arthritis in mice. J Immunol 184, 379–390, doi:10.4049/jimmunol.0901639 (2010).

35. Nandakumar, K. S., Svensson, L. & Holmdahl, R. Collagen type II-specific monoclonal antibody-induced arthritis in mice: description of the disease and the influence of age, sex, and genes. Am J Pathol 163, 1827–1837, doi:10.1016/S0002-9440(10)63542-0 (2003).

36. Neeli, I., Khan, S. N. & Radic, M. Histone deimination as a response to inflammatory stimuli in neutrophils. J Immunol 180, 1895–1902, doi:10.4049/jimmunol.180.3.1895 (2008).

37. Khandpur, R. et al. NETs are a source of citrullinated autoantigens and stimulate inflammatory responses in rheumatoid arthritis. Sci Transl Med 5, 178ra140, doi:10.1126/scitranslmed.3005580 (2013).

38. de Bont, C. M. et al. Autoantibodies to neutrophil extracellular traps represent a potential serological biomarker in rheumatoid arthritis. J Autoimmun 113, 102484, doi:10.1016/j.jaut.2020.102484 (2020).

39. Brinkmann, V. et al. Neutrophil extracellular traps kill bacteria. Science 303, 1532–1535, doi:10.1126/science.1092385 (2004).

40. Berthelot, J. M., Le Goff, B., Neel, A., Maugars, Y. & Hamidou, M. NETosis: At the crossroads of rheumatoid arthritis, lupus, and vasculitis. Joint Bone Spine 84, 255–262, doi:10.1016/j.jbspin.2016.05.013 (2017).

41. Perez-Sanchez, C. et al. Diagnostic potential of NETosis-derived products for disease activity, atherosclerosis and therapeutic effectiveness in Rheumatoid Arthritis patients. J Autoimmun 82, 31–40, doi:10.1016/j.jaut.2017.04.007 (2017).

42. Chang, M. H. & Nigrovic, P. A. Antibody-dependent and -independent mechanisms of inflammatory arthritis. JCI Insight 4, doi:10.1172/jci.insight.125278 (2019).

43. Mayadas, T. N., Tsokos, G. C. & Tsuboi, N. Mechanisms of immune complex-mediated neutrophil recruitment and tissue injury. Circulation 120, 2012–2024, doi:10.1161/CIRCULATIONAHA.108.771170 (2009).

44. Lux, A., Yu, X., Scanlan, C. N. & Nimmerjahn, F. Impact of immune complex size and glycosylation on IgG binding to human FcgammaRs. J Immunol 190, 4315–4323, doi:10.4049/jimmunol.1200501 (2013).

45. Rojko, J. L. et al. Formation, clearance, deposition, pathogenicity, and identification of biopharmaceutical-related immune complexes: review and case studies. Toxicol Pathol 42, 725–764, doi:10.1177/0192623314526475 (2014).

46. Pawankar, R., Holgate, S. T. & Rosenwasser, L. J. Allergy frontiers. (Springer, 2009).

47. Denning, N. L., Aziz, M., Gurien, S. D. & Wang, P. DAMPs and NETs in Sepsis. Front Immunol 10, 2536, doi:10.3389/fimmu.2019.02536 (2019).

48. Sokolove, J., Zhao, X., Chandra, P. E. & Robinson, W. H. Immune complexes containing citrullinated fibrinogen costimulate macrophages via Toll-like receptor 4 and Fcgamma receptor. Arthritis Rheum 63, 53–62, doi:10.1002/art.30081 (2011).

49. Tsourouktsoglou, T. D. et al. Histones, DNA, and Citrullination Promote Neutrophil Extracellular Trap Inflammation by Regulating the Localization and Activation of TLR4. Cell Rep 31, 107602, doi:10.1016/j.celrep.2020.107602 (2020).

50. Paues Goranson, S., et al. Circulating H3Cit is elevated in a human model of endotoxemia and can be detected bound to microvesicles. Sci Rep 8, 12641, doi:10.1038/s41598-018-31013-4 (2018).

51. Tian, Y. et al. Serum citrullinated histone H3 concentrations differentiate patients with septic verses non-septic shock and correlate with disease severity. Infection 49, 83–93, doi:10.1007/s15010-020-01528-y (2021).

52. Li, Y. et al. Citrullinated histone H3: a novel target for the treatment of sepsis. Surgery 156, 229–234, doi:10.1016/j.surg.2014.04.009 (2014).

53. Deng, Q. et al. Citrullinated Histone H3 as a Therapeutic Target for Endotoxic Shock in Mice. Front Immunol 10, 2957, doi:10.3389/fimmu.2019.02957 (2019).

54. Steen, J. et al. Recognition of Amino Acid Motifs, Rather Than Specific Proteins, by Human Plasma Cell-Derived Monoclonal Antibodies to Posttranslationally Modified Proteins in Rheumatoid Arthritis. Arthritis Rheumatol 71, 196–209, doi:10.1002/art.40699 (2019).

55. Reijm, S. et al. Cross-reactivity of IgM anti-modified protein antibodies in rheumatoid arthritis despite limited mutational load. Arthritis Res Ther 23, 230, doi:10.1186/s13075-021-02609-5 (2021).

56. Mannoor, K., Matejuk, A., Xu, Y., Beardall, M. & Chen, C. Expression of natural autoantibodies in MRL-lpr mice protects from lupus nephritis and improves survival. J Immunol 188, 3628–3638, doi:10.4049/jimmunol.1102859 (2012).

57. Werwitzke, S. et al. Inhibition of lupus disease by anti-double-stranded DNA antibodies of the IgM isotype in the (NZB x NZW)F1 mouse. Arthritis Rheum 52, 3629–3638, doi:10.1002/art.21379 (2005).

58. Brewer, R. C. et al. Oral mucosal breaks trigger anti-citrullinated bacterial and human protein antibody responses in rheumatoid arthritis. Sci Transl Med 15, eabq8476, doi:10.1126/scitranslmed.abq8476 (2023).

59. Li, S. et al. Autoantibodies From Single Circulating Plasmablasts React With Citrullinated Antigens and Porphyromonas gingivalis in Rheumatoid Arthritis. Arthritis Rheumatol 68, 614–626, doi:10.1002/art.39455 (2016).

